# Inducible degradation of the *Drosophila* Mediator subunit Med19 reveals its role in regulating developmental but not constitutively-expressed genes

**DOI:** 10.1101/2022.06.06.494962

**Authors:** Denis Jullien, Emmanuelle Guillou, Sandra Bernat-Fabre, Adeline Payet, Henri-Marc G. Bourbon, Muriel Boube

## Abstract

The multi-subunit Mediator complex plays a critical role in gene expression by bridging enhancer-bound transcription factors and the RNA polymerase II machinery. Although experimental case studies suggest differential roles of Mediator subunits, a comprehensive view of the specific set of genes regulated by individual subunits in a developing tissue is still missing. Here we address this fundamental question by focusing on the Med19 subunit and using the *Drosophila* wing imaginal disc as a developmental model. By coupling auxin-inducible degradation of endogenous Med19 *in vivo* with RNA-seq, we got access to the early consequences of Med19 elimination on gene expression. Differential gene expression analysis reveals that Med19 is not globally required for mRNA transcription but specifically regulates positively or negatively less than a quarter of the expressed genes. By crossing our transcriptomic data with those of *Drosophila* gene expression profile database, we found that Med19-dependent genes are highly enriched with spatially-regulated genes while the expression of most constitutively expressed genes is not affected upon Med19 loss. Whereas globally downregulation does not exceed upregulation, we identified a functional class of genes encoding spatially-regulated transcription factors, and more generally developmental regulators, responding unidirectionally to Med19 loss with an expression collapse. Moreover, we show *in vivo* that the Notch-responsive *wingless* and the *E(spl)-C* genes require Med19 for their expression. Combined with experimental evidences suggesting that Med19 could function as a direct transcriptional effector of Notch signaling, our data support a model in which Med19 plays a critical role in the transcriptional activation of developmental genes in response to cell signaling pathways.

**Author summary:** The Mediator is a large evolutionarily conserved multisubunit complex that plays essential functions in gene expression by relaying cues emanating from enhancer-bound transcription factors to the RNA polymerase II machinery. The transcriptional landscapes regulated by each Mediator subunit, especially *in vivo* in the context of developmental processes, remains poorly characterized. We therefore sought to provide a comprehensive view of the genes directly regulated by Med19, an archetypal Mediator subunit, in the context of the development of the *Drosophila* wing. Our work has important methodological implications as to carry out our analysis in the best experimental conditions, we generated mutant flies in which we could trigger fast degradation of the Med19 subunit on demand in the tissues of the living animal. We found that only a small part of the genes expressed in the developing wing are controlled by Med19 showing that this Mediator component exercises specific functions. Another major finding is the strong involvement of Med19 in the regulation of the expression of the genes involved in developmental processes supporting the idea that Med19 is a Mediator component dedicated to highly regulated mode of gene expression. The most unanticipated discovery of our study is the fact that Med19 does not appear to play a role in the expression of the genes that are constitutively transcribed, among which the housekeeping genes. This raises the notion that in the context of simple —non-regulated— mode of expression, part of the Mediator functionalities, among which Med19, otherwise dedicated to highly regulated type of transcription, may become dispensable.

## Introduction

Mediator is an eukaryotic multisubunit complex [1, 2] that plays multiple functions in the regulation of RNA Polymerase II (RNA PolII) [3, 4]. In metazoans, it comprises 30 different subunits together acting as functional bridge between enhancer-bound transcription factors (TFs) and the RNA PolII machinery associated with core promoter elements [5]. A current challenge in understanding the Mediator functions is to unravel the role of its individual subunits. Indeed, although the complex has been shown to behave globally as an essential RNA PolII cofactor [6–8], several Mediator subunits have been shown to carry out specific functions as revealed by genetic data [9]. This notion is exemplified by a differential requirement for cell viability as observed in genetic studies made in *Drosophila* [10, 11], as well as in mammalian cells [6]. Yet the precise nature of the genes regulated by each Mediator subunit remains poorly characterized, particularly in the context of the development where fine tuning of gene expression is critical and during which genetic studies show that most of the Mediator subunits play an essential role. This lack of data is partly due to the fact that assessing the direct consequences on gene expression of depleting a given Mediator subunit in a developing tissue is hampered first by the difficulty to obtain such tissue due to deleterious effects of losing the function of a Mediator component, and second by the problem of secondary transcriptional cascade effects produced by extended loss of function periods. Recent studies have shown the great interest of conditional protein degradation systems [12], where induction of fast degradation of a Mediator subunit allows to access the primary consequences on gene expression [6–8,13].

Here we addressed the fundamental question of the early consequences of losing the function of a Mediator component on gene expression using the *Drosophila* larval wing imaginal disc as a developmental model. We focused on Med19, a key Mediator subunit previously shown to interact with several TFs (GATA, HOX and REST) [14–16] and the RNA PolII but whose transcriptional landscape it regulates still remains to be characterized. We introduced the sequence encoding the Auxin Induced Degron (AID) [17] into the *Drosophila Med19* gene generating an allele encoding a degradable version of the protein that achieved inducible, fast, and full depletion in the animal. By combining this unique tool with RNA sequencing of the transcripts of the wing imaginal disc, we observed that less than 25% of the genes expressed in the wing disc were deregulated early on after Med19 depletion. Strikingly, among the genes whose expression was not affected were most of the constitutively expressed genes. Conversely, we found that the genes deregulated following Med19 degradation were highly enriched in those exhibiting specific spatio-temporal expression patterns. Moreover, the finding that the transcription of nearly all spatially-regulated transcription factors, and notably key Notch responsive genes, requires *Med19* function points to the idea that Med19 plays a specialized role in promoting the expression of developmental regulators.

## Materials and methods

### DNA constructs cloning

The DNA sequence of the guide RNA – CTTATGTCGCAGTTTTAGGC – obtained by annealing two complementary DNA oligonucleotides: M19gRNA Fw and M19gRNA Rv, was cloned to the *Bbs*I site of the pCFD3 vector [18], according to the protocol provided by CRISPRfly design (http://www.crisprflydesign.org), resulting in the pCFD3 M19gRNA plasmid. The donor construct (Fig. 1B), carried in a pUC19 backbone, was generated by the assembly of 5 DNA fragments using the NEBuilder HiFi DNA assembly cloning kit (New England Biolabs). These DNA fragments, namely pUC19, the 5’ homology arm, AID, GFP, and the 3’ homology arm were produced by PCR (Phusion DNA polymerase, New England Biolabs). The 5’ and 3’ homology arm DNA fragments were amplified from genomic DNA prepared from *vasa*-Cas9 flies. The source of the AID degron DNA sequence was the pAID- AsiSI plasmid provided by the G. Legube lab (CBI, Toulouse, France). The sequence of the primers used for constructs cloning are available in S1 table.

**Fig 1.**
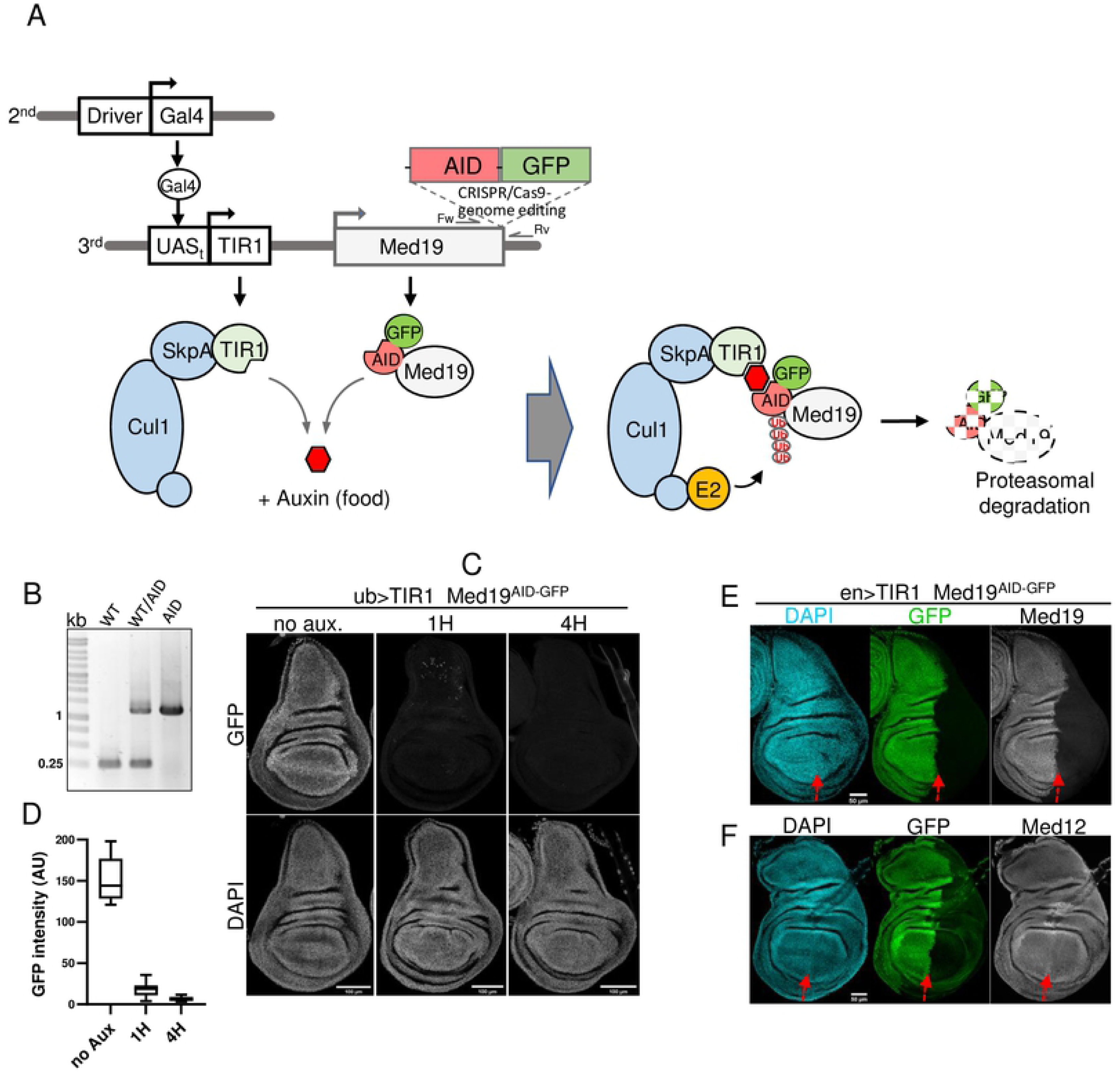
K-in of the Auxin Induced Degron (AID) for fast and deep depletion of Med19 in the *Drosophila* wing imaginal disc. (A) Principle of auxin-dependent degradation of Med19^AID^ in *Drosophila*. The Med19 endogenous gene located on the third chromosome has been engineered using CRISPR-Cas9 system in order to generate Med19 fusion protein with AID and GFP tags. The expression of the auxin-dependent and AID-specific F-box protein TIR1 is controlled by the UAS/Gal4 system. The Gal4 driver is located in the second chromosome, and the *UAS-TIR1* construct in the third chromosome. The F-box TIR1 incorporates the endogenous SCF (Skip1 Cul1 F-box) E3 ubiquitin ligase complex. Auxin (added to the fly food) triggers poly-ubiquitination of Med19^AID^ by the SCF-TIR1 targeting the Mediator subunit to degradation by the proteasome. (B) PCR genotyping for the insertion of the AID-GFP coding sequence in the *Med19* locus using the primers couple indicated in (A) with genomic DNA extracted from WT control (Med19^+^ left lane), Med19^+^/*Med19^AID^*heterozygous (middle lane), or *Med19^AID^* homozygous (right lane) adult flies. WT and *Med19^AID^* alleles theoretically produce a 270 and 1200 pb PCR product, respectively. (C) Ubiquitous Auxin-dependent degradation of Med19^AID^ in the wing imaginal disc. Images showing DAPI and GFP (detection of Med19^AID^) signals from *ub>TIR1; Med19^AID^* wing discs dissected from larvae not exposed to auxin (no aux.), or fed with auxin for 1 hour (1H), or 4 hours (4H). (D) GFP signal quantification (Arbitrary Unit), as measurement of Med19^AID^ depletion, in *ub>TIR1; Med19^AID^* wing imaginal discs (images shown in (C)) obtained from larvae not exposed to auxin (no Aux., n=9), or fed with auxin during 1 hour (1H, n=11), or 4 hours (4H, n=12). (E) Auxin dependent degradation results in loss of both C-terminal GFP and N-terminal Med19 moieties of Med19^AID^. Images showing *en>TIR1 Med19^AID^*wing discs, in which auxin-induced degradation in the posterior compartment was performed for 4 hours, co- immunostained using anti-GFP and Med19 antibodies. (F) Auxin dependent degradation of Med19^AID^ does not affect Med12 expression. Co-immunostaining as in (E) except antibodies against Med12 were used.

### CRISPR/Cas9 mediated Knock-in of AID-GFP to the *Med19* locus

Four hundred *Vasa*-Cas9 (BDSC #51323) embryos were injected with a mixture of the donor plasmid and the pCFD3 M19gRNA. The resulting mosaic adults were crossed with double balanced *TM3*/*TM6B* flies. The F1 progeny was individually crossed to *TM3*/*TM6B* flies before being PCR-genotyped for the insertion of the AID-GFP cassette using the M19genotFw and M19genotRv primer couple (S1 Table). The progeny of a fly that came out positive from the genotyping were crossed together to produce a homozygous stock for the degradable Med19- AID-GFP allele (referred as to *Med19^AID^*).

### *Drosophila* stocks and crosses

Flies were raised at 25°C on standard yeast agar wheatmeal medium. V*as*a-Cas9, *UAS-TIR1* (#76123), the following Gal4 drivers (annotated with “>”) located on the 2^nd^ chromosome: *ubiquitin* (#32551), *engrailed* (#30564), *apterous* (#3041) were obtained from Bloomington BDSC. To associate the *Med19^AID^* allele with the expression of the F-box protein TIR1 under the control of a Gal4 driver, *Med19^AID^*and *UAS-TIR1* were recombined. The recombined 3^rd^ chromosome was then genetically associated with a 2^nd^ chromosome bearing either *ubiquitin* (*ub*>), *engrailed* (*en*>) or *apterous* (*ap*>) Gal4 driver. The lines thus generated *ub*>/*CyO*; *Med19^AID^ UAS-TIR1*, *en*>; *Med19^AID^ UAS-TIR1*, *ap*>/*CyO*; *Med19^AID^ UAS-TIR1* were used to degrade Med19^AID^ in all the cells, the posterior compartment, the dorsal domain of the wing disc, respectively.

To trigger Med19^AID^ degradation, L3 larvae were transferred to a 30 mm dish filled with 10 ml of a solid medium containing 0.8% bacteriological agar, 4% sucrose, and 2.5 mM of the auxin analog NAA (1-Naphtaleneacetic acid, Sigma N0640). The dish was then sealed with a piece of parafilm. To ensure gas exchange, small holes were made to the seal with a needle, and the dish was incubated at 25°C for required amount of time.

### RNA-seq differential gene expression analysis

80 wing discs were hand-dissected from female *ub>Med19^AID^*L3 larvae either fed with food containing NAA for a duration of 4 hours (UbAux), or kept in the same food devoid of NAA (Ub). RNA extraction was performed using TRIzol reagent (Thermofisher). The process was repeated as biological triplicate. Poly-A+ RNA was purified and paired end sequenced on a NovaSeq 6000 (Illumina). The bioinformatic analysis was performed using locally installed Galaxy instance (http://sigenae-workbench.toulouse.inra.fr). The quality control of sequences was done with FastQC (v0.11.4), then reads were aligned using TopHat2 (v2.0) on the *Drosophila melanogaster* dm6 genome assembly. Paired reads were counted by gene using htseq (v1.0). These counts were used to perform differential gene expression analyses using DESeq2 (1.26.0) in R comparing UbAux with Ub. Gene Ontology (GO) enrichment analysis (biological process) of the genes exhibiting an expression Log2FC <-1 or >1 was performed using ShinyGO [19] http://bioinformatics.sdstate.edu/go/.

### Gene expression profile analyses

Expression scores in 58 different *Drosophila* tissues and developmental stages (detailed S2 Table) of each genes found significantly expressed in the wing imaginal disc were extracted from the modENCODE *Drosophila* RNA expression profiling project using DGET tool [20] https://www.flyrnai.org/tools/dget/web/. The HK* and SR* gene lists (S4 Table) analyzed in the Figure 3 C–F were generated as followed. To select for constitutively expressed genes – the HK* genes – using the expression scores matrix, those containing any of 0, 1, 2, 3 score (null or very low expression) in their expression profile were excluded. For SR* genes, the selection criteria were 0 score occurrence ≥ 7 and sum of occurrence of 0, 1, and 2 scores ≤ 52. Heat maps, box plots, violin plots, and statistical analysis were made using Prism9 (Graphpad). The maximal expression value in the color range of the heat maps was set to 10 for a better visualization of the difference of expression profile.

### RNA quantification by RT-qPCR

RNA, trizol extracted from wing discs of L3 larvae, were reverse transcribed using SuperScriptTM II Reverse Transcriptase (Thermo Fisher Scientific) and cDNA were quantified by real-Time qPCR (CFX Bio-Rad) with specific oligonucleotides (S1 Table). Absolute quantification of each mRNA was normalized to the mean expression of Actin 42A mRNA in the same sample. mRNA measured in tissue from female *ub> Med19^AID^* L3 larvae fed without NAA (ub), was set at 100% to compare with the ones fed 4h with NAA (ub+auxin).

### Immunostainings

L3 wing imaginal discs attached to inverted larvae were fixed in PBS 4% paraformaldehyde for 20 minutes, and permeabilized by incubation in PBS 0.1% Triton X100 for 30 minutes. Primary antibodies diluted in PBS 1% BSA were applied overnight at 4°C. Incubation with Alexa Fluor-conjugated secondary antibodies was carried out at 22°C for 2 hours, and DNA was stained with DAPI (4’,6-diamidino-2-phenylindole) while washing the secondaries. Wing discs were then manually dissociated from carcasses, and mounted in Vectashield mounting medium. The following antibodies were used: guinea pig anti-Med19 1:1000 [14], mouse anti-wingless 1:100 (4D4; DSHB), mouse anti-achaete (1:100, DSHB), rabbit anti-GFP (1:1000).

Images were acquired using Zeiss LSM710 (20X PL APO ON 0,8; 40x PL APO oil DIC ON 1,3), or Leica SP8 (20x multi immersion PL APO ON 0,75; 40x HC PL APO oil ON 1,30) confocal microscopes and were analyzed and quantified using Fiji/ImageJ.

### smiFISH

For RNA FISH, smiFISH [21] was performed according to the *Drosophila* wing discs protocol available from https://www.biosearchtech.com/support/resources/stellaris-protocols with the following modifications. Dissection and fixation were carried out as for the immunostaining except that the fixation time was 26 minutes, after what fixed material was washed 3 times in PBS to remove the paraformaldehyde, and then kept in 70% ethanol at 4°C for up to one week. DNA oligonucleotide primary probes targeting introns for *wingless* and *achaete*, the coding sequence for – intronless genes – E(spl)m3 and E(spl)mbeta were designed using the online Stellaris probe designer tool (https://www.biosearchtech.com/stellaris-designer). Primary probes were labeled using a Cy3 conjugated FlapX DNA oligonucleotide. All the DNA oligonucleotides used in smiFISH were obtained from Integrated DNA Technologies.

### *In vitro* protein binding assays

GST (produced from empty pGex4T) and GST::Med19 proteins were expressed in BL21 AI *E. coli* (37°C, 1 H, with 500µM IPTG in LB media), from which soluble forms were purified on glutathione-Sepharose beads (GE Healthcare). Recombinant C-terminally HA tagged Su(H) was produced using *in vitro* coupled transcription/translation (TNT with rabbit reticulocyte extracts; Promega Inc.) from DNA matrices generated by PCR using Su(H)-HA forward and reverse primers combined with the *Su(H)* cDNA (obtained from the *Drosophila* Genomic Resource Center (DGRC) Gold Collection) as template. GST pulldown was performed as previously described [22]. The following antibodies were used for western blotting: mouse anti-HA (1:1,000; Covance). Secondary HRP conjugated antibodies were Goat anti mouse-HRP (1:10000).

## Results

### Inducible degradation of the Mediator subunit Med19 in *Drosophila* larvae

In order to achieve an inducible and fast depletion of Med19 *in vivo* in *Drosophila* wing imaginal disc upstream of RNA-seq transcriptional profiling, we turned to the Auxin Induced Degron (AID) [17] conditional protein degradation system that has been shown to be particularly well suited to this model organism (Fig.1A) [23–25]. We inserted the coding sequence of the AID fused to GFP into the endogenous *Med19* locus, using CRISPR-cas9 genome editing (S1 Fig.), C-terminally tagging Med19. Proper insertion of the AID-GFP cassette was validated by PCR genotyping (Figure 1B). Importantly, flies homozygous for this degradable and fluorescent *Med19* allele, hereafter referred to as *Med19^AID^*, were viable, fertile and did not exhibit any visible phenotype, indicating that the addition of the AID-GFP moiety does not affect Med19 function. We next generated a genetic setup in which UAS/Gal4-based expression of the AID cofactor TIR1 controls ubiquitous or regionalized degradation of Med19^AID^ depending on the type of Gal4 driver used (Fig.1A). Central to the feasibility of our study was the possibility to carry out efficient auxin-dependent depletion of Med19^AID^ in all the cells of the wing imaginal disc. We found this could be achieved using the ubiquitin driver (*ub*>), as feeding *ub>TIR1 Med19^AID^* larvae with auxin (Fig. 2A) resulted in fast degradation of MED19^AID^, with less than 10% remnant levels after one hour, and almost complete disappearance in nearly all the nuclei of the wing disc after four hours (Figure 1C-D). Similar auxin-induced degradation speed and depth was observed with the *en>TIR1 Med19^AID^* setup in which *engrailed*Gal4 drives depletion in the posterior compartment of the wing disc (S2 Fig.). Using this regionalized degradation, we confirmed that the Med19 moiety of the MED19^AID^ fusion protein was actually degraded along with the C-terminal GFP by immunostaining using antibodies against Med19 (Figure 1E). In addition, we observed that the level of Med12, another Mediator subunit, was not affected upon MED19^AID^ degradation (Figure 1F) providing additional evidence for the specificity of the depletion. These data therefore demonstrate that we developed an AID-based Med19 degradation system performing fast and efficient depletion of this Mediator subunit *in vivo*, thereby opening the possibility of investigating the early consequence of Med19 loss in gene expression in the developing wing disc.

**Fig 2.**
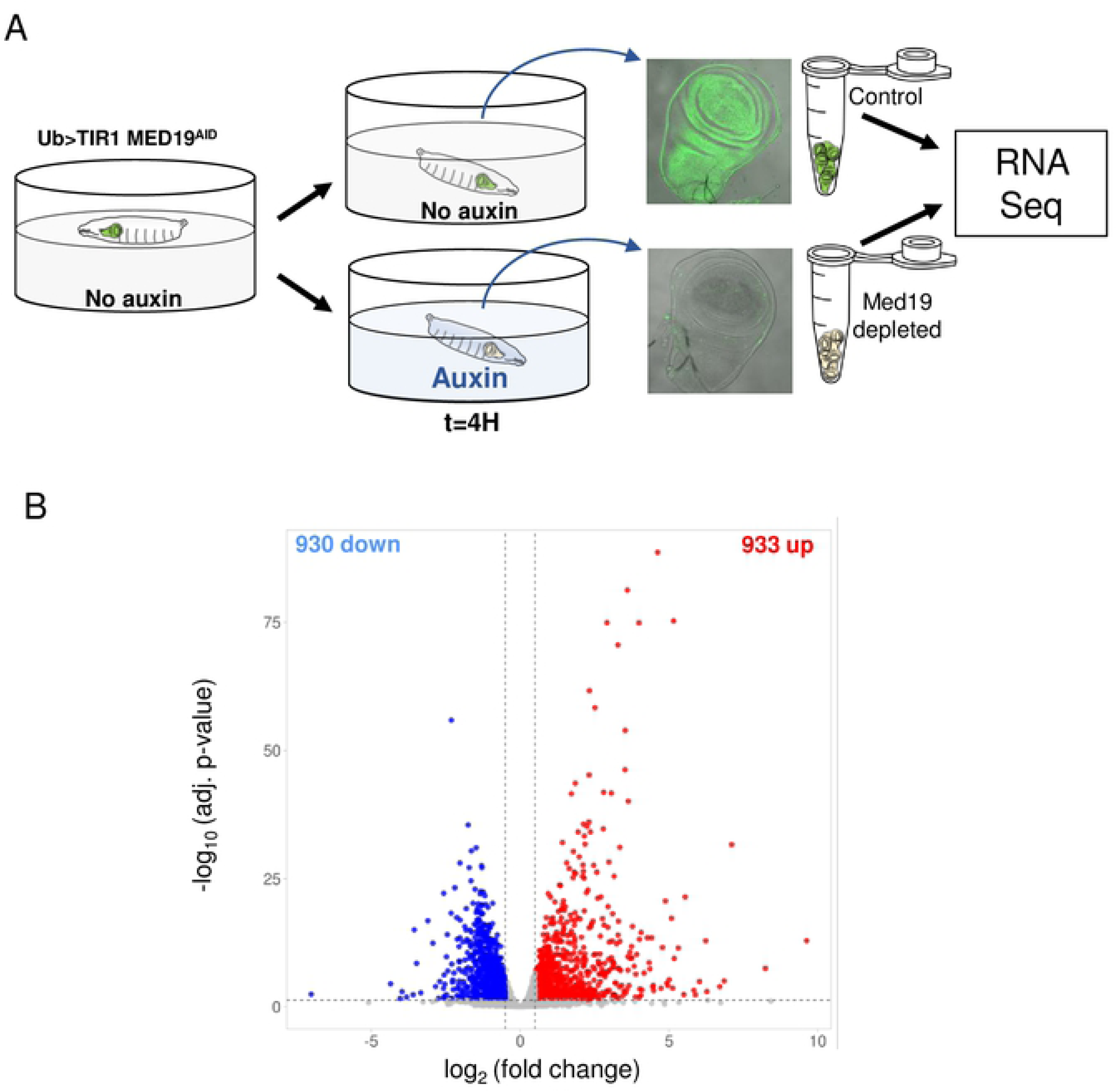
Med19^AID^ removal results in the deregulation a limited part of the wing imaginal disc transcriptome. (A) Scheme depicting the experimental setup used to produce wing imaginal discs in which Med19AID degradation has been triggered, or not (no Auxin control), upstream of the RNA-seq analysis. (B) Differential gene expression analysis volcano plot showing the expression fold change after Med19^AID^ degradation and associated adjusted p-values. 8753 genes found expressed in the wing imaginal disc are depicted. Genes with significative fold change (adjusted p-value <0,05) are shown in blue for the down-regulated (log2FC <-0,5) and in red for the up-regulated (log2FC >0,5).

### Med19 removal results in the deregulation of only a subset of the genes expressed in the wing imaginal disc

Having setup an unprecedented tool to trigger Med19 depletion in the wing imaginal disc, we next proceeded to the analysis of the early consequences of losing Med19 function on the transcriptome. We prepared RNA samples from wing discs dissected from larvae that were fed for four hours with a medium containing or not auxin (Fig. 2A), proceeded to their sequencing, and analyzed the data for differential gene expression (DGE). Globally, we found that the expression of most *Drosophila* genes (78%) was unaffected upon Med19^AID^ degradation. Among the 1863 genes showing significant changes in expression (adjusted p-value <0.05), we found a similar proportion of genes downregulated or upregulated (Figure 2B). Although the Mediator complex is commonly seen as indispensable for RNA PolII-dependent transcription, these data suggested that the *Drosophila* Med19 subunit is instead specifically required for the proper activation or repression of a limited number of genes. Consistent with this observation, we previously showed that the loss of Med19 is not strictly required for cell viability [14] implicating that expression of the housekeeping genes does not depend on Med19. Indeed, we found upon a preliminary survey of the DGE data that the expression of several families of canonical housekeeping (HK) genes (*e.g.,* encoding ribosomal proteins, Mediator subunits, general transcription factors (GTFs) or RNA PolII subunits, list in S3 Table) was largely unchanged after Med19^AID^ removal (S3 Fig.). These results raised the idea that Med19 could be more generally dispensable for the expression of constitutively expressed genes and prompted us to examine whether constitutive or highly regulated expression mode determine the sensitivity of the transcription of a gene to Med19 removal.

### Med19 depletion affects the expression of spatially restricted but not constitutively expressed genes

We looked for a link between the expression change upon Med19^AID^ loss and the expression mode (constitutive versus highly regulated). We extracted from the modENCODE *Drosophila* gene expression database the expression profiles of each gene we found expressed in the wing imaginal disc, compiling expression scores in 58 different developmental stages and tissues (Fig3A left panel and table 2 for details) –[26]. We next used the resulting expression scores matrix to generate heat maps in which the expression profile of each gene (columns in the heat maps shown in Fig. 3) was sorted as a function of the value of the expression fold change (horizontal axis Fig. 3A). Strikingly, comparing the expression profile heat maps of the genes whose expression is the most affected (Log2FC>1 and <-1) with that of those the less impacted (-0.03<Log2FC<0.03) revealed that while the expression patterns of the up- and down- regulated Med19 target genes are similar, those of the unaffected genes are clearly different from the two others. Specifically, in down- and up-regulated genes, null expression occurrence (dark blue color code) appeared frequently (Fig. 3A left and right heat maps), whereas it was clearly much sparse in genes whose expression showed little variation (Fig. 3A Log2FC around 0, middle panel). We statistically validated this bias by counting the number of null expression score occurrences in the expression profile of each gene affected (|Log2FC|>1) and unaffected (-0.1<Log2FC<0.1) in Med19AID loss. This analysis revealed a significant difference between the count means (P<0.0001), the number of null expression count in non-deregulated genes being much lower than in those deregulated (Fig. 3B). By definition genes spatially and temporally regulated – further called spatially restricted genes (SR) - contain in their developmental and tissue expression profile null expression occurrences when conversely absence of null expression is a hallmark of ubiquitously expressed HK genes. Therefore, these results indicated that SR genes were enriched among those affected by Med19 depletion.

**Fig 3.**
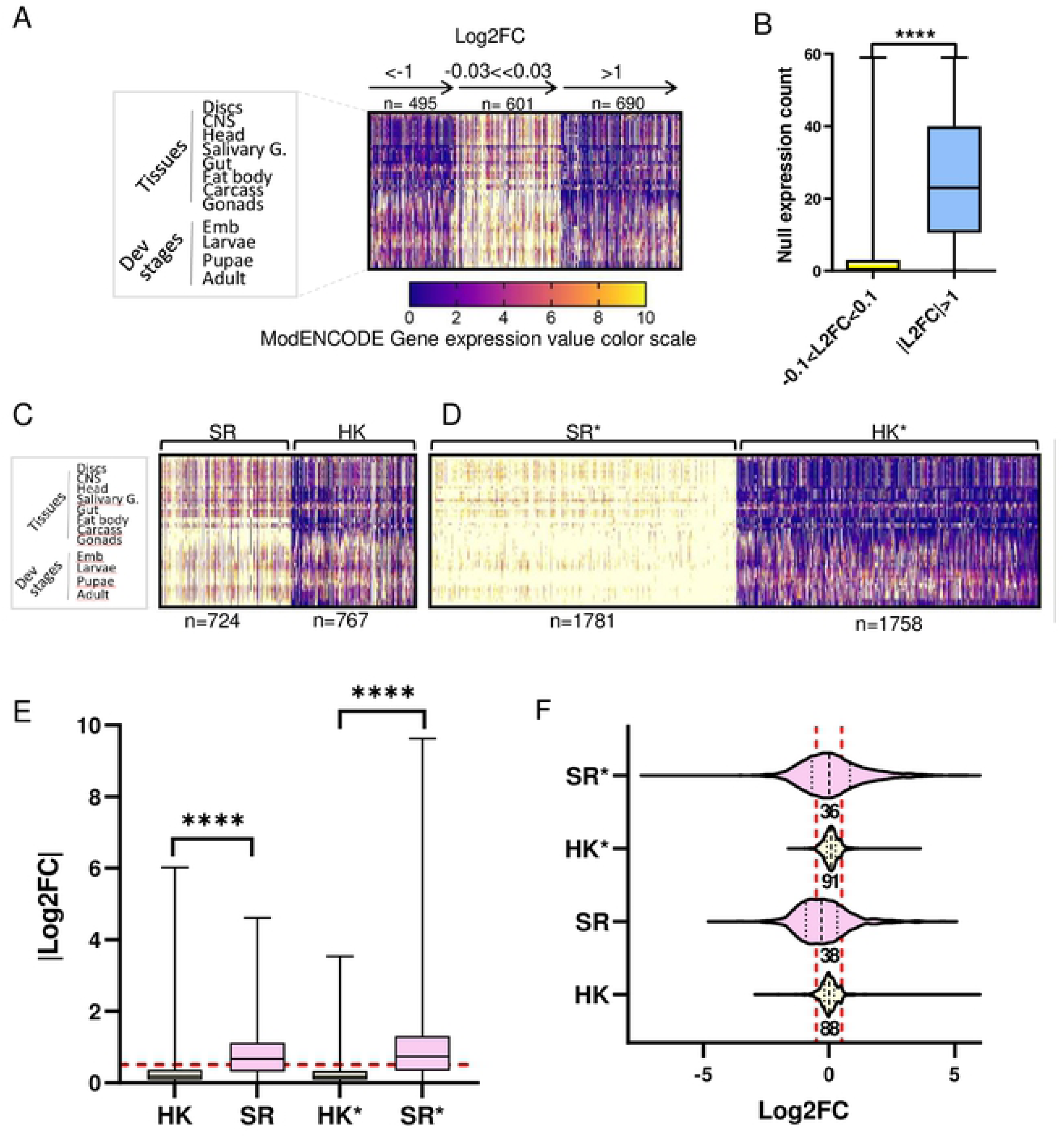
Spatially and temporally regulated genes are more prone to expression change than constitutively expressed genes upon Med19^AID^ depletion. (A) Heat map depicting the spatio temporal expression profiles (modENCODE) of genes belonging to three different ranges of expression fold change (L2FC specified above the horizontal axis) in our transcriptomic analysis. Each column contains the color-coded expression score (modENCODE) of a gene in 58 different tissues and developmental stages (left panel and table S2 for details) extracted from the modENCODE *Drosophila* gene expression database the expression profiles of each gene we found expressed in the wing imaginal disc, compiling expression scores in 58 different developmental stages and tissues. (B) Box plot analyzing the count of the number of null expression (0 and 1 expression score occurrences) among the 58 expression values for the genes belonging to |L2FC|>1 (n=1185) or -0.1<L2FC<0.1 (n=1704) range. Median significantly different P<0.0001 two-tailed non-parametric Mann Whitney test. (C) Gene expression profile heat maps as in (A) comparing the expression profiles of published sets of housekeeping genes (HK, n=767) and spatially regulated genes (SR, n=724); lists in S4 Table. (D) Heat maps as in (C) showing the expression profiles of the HK* and SR* sets of genes obtained by selecting HK and SR-like expression profiles from the modENCODE expression scores matrix (see materials and methods, lists in S4 Table). (E) Box plot comparing the Log2FC absolute values of the sets of genes specified on the x axis, and whose expression profile is shown above. ****: median significantly different P<0.0001 two-tailed non-parametric Mann Whitney test. The red dashed line marks |Log2FC|= 0.5. (F) Violin plot showing the distribution of the expression Log2FC values of the sets of genes mentioned on the vertical axis (analyzed in the figure 2 E). The percentage of genes in the -0.5<Log2FC<0.5 range, delimited by the two red dashed vertical lines, is indicated below the violin plot.

To further explore this notion, we performed the reverse approach by examining the expression fold change values upon Med19 depletion in lists of HK and SR genes obtained either from published studies or that we established from the expression profile matrix of the genes transcribed in the wing imaginal disc. The published sets of genes (S4 Table) consist of 767 HK genes identified in a large-scale *in situ* hybridization project as ubiquitously expressed throughout *Drosophila* embryogenesis [27, 28], and a list of 724 SR genes derived from a single-cell RNA-seq study made in the wing imaginal disc [29]. In addition, we sought to expand our analysis to other sets of HK and SR genes obtained by an independent approach. By applying simple selection criteria to the modENCODE gene expression matrix, based on the presence or absence of null expression score (selection details in the Material and Methods), we generated large sets of genes (total of 3538 genes) exhibiting either HK- or SR-like expression profile (Fig. 3, Heat maps in C to be compared with D) hereafter referred to as HK* and SR* (S4 Table). Of note, almost half of the HK and SR genes were contained in the larger HK* and SR* gene lists, respectively (S4 Fig.), validating the mode of selection of the latter. Comparing the values of the expression fold change upon Med19^AID^ depletion between HK and SR as well as between HK* and SR* genes revealed that this indicator was significantly higher in spatially restricted genes than in the housekeeping genes in both selection modes (Fig. 3E). In addition, examination of the distribution of the Log2FC values showed that 88% of the HK genes and 91% of the HK* genes fell in the FC range (-0.5<Log2FC<0.5) where gene expression is considered not to be affected by Med19^AID^ depletion while this proportion dropped to 38% and 36% with the SR and the SR* genes, respectively (Fig. 3F). Therefore, the analysis of a significant part of the HK and SR genes expressed in the wing imaginal disc (1884 and 1822 genes, respectively) indicates that most of the HK genes are not affected by Med19 depletion, whereas the expression of an important proportion of the SR genes, but not all of them, depends on Med19 activity. In conclusion, the fact that Med19 loss of function disrupt or not the expression of a gene seems to depend on its mode of expression. Yet this simple binary model is likely an oversimplification of the reality given that about a third of the SR genes appears insensitive to Med19 degradation.

After investigating the relationship between the mode of expression and the deregulation of the expression, we next focused on the function of the genes affected by *Med19* loss of function.

### Med19 promotes the expression of developmental regulators

Transcription factors (TFs) whose expression is highly regulated spatio-temporally are functionally associated with development, and are commonly considered as developmental regulators. We analyzed the distribution of the value of the expression FC of the TF-encoding genes expressed in the wing imaginal disc (Fig. 4A), and focused on those with a documented evidence [29] for a spatially-restricted (SR) expression pattern (Fig. 4A, purple colored) in the wing disc (SR TFs). Unexpectedly, we found a strong bias to the downregulation regarding their distribution (Fig. 4A). More precisely, plotting the proportion of SR TFs in each category of expression change (Fig. 4B) shows that 85% of the SR TFs are deregulated upon Med19^AID^ degradation, in line with our previous conclusion concerning the SR genes. However, contrary to the general behavior in which an equal distribution between down and up regulation is seen (Fig. 2B), the vast majority (91%) of the SR TFs deregulated upon Med19^AID^ loss fall in the group of the downregulated. We experimentally validated this finding by measuring using RT-qPCR the transcripts level of three developmental TFs (encoding by *cut*, *salm* and *E(spl)m8*),(Fig. 4C) and imaging the expression of the TF-encoding *achaete* (*ac*) gene following wing disc immunostaining (S5A Fig.). In agreement with our conclusions, we observed that the expression of these four developmental TFs in the wing imaginal disc relies on Med19 activity (Fig. 4C, and S5A Fig.), whereas the transcripts level of two HK genes (eEF1, and myosin) do not significantly change after Med19^AID^ depletion (Fig4C). We next sought to expand our analysis to a developmental gene whose product is not a TF: *wingless* (*wg*) encoding the ligand of the Wnt signaling, known to be a key regulator of wing development. RT-qPCR and wing disc immunostaining both indicated a reduction in *wg* expression upon Med19^AID^ removal (Fig 4C and S5B Fig.). To examine whether the effect of Med19^AID^ depletion on *wg* expression is transcriptional, the short lived unspliced *wg* pre-mRNAs were visualized by RNA FISH using intronic probes. Using this approach, we observed that the level of *wg* pre-messengers was markedly reduced specifically in the wing disc territory where Med19^AID^ depletion occurred (Fig. 4C-D), indicating that Med19 is required for *wg* transcription. Moreover, the fact that *wg* expression is already affected one hour after Med19^AID^ degradation initiation (Figure 4D) suggests a direct coupling between Med19 function and *wg* transcription. Therefore, our results revealed the existence of sets of SR genes, functionally related to the development, for which, strikingly, Med19 acts exclusively as a positive transcriptional factor. To further explore this idea with a more global approach, we analyzed the Gene Ontology enrichments of the genes down-regulated after Med19^AID^ loss. Among the 30 top biological processes identified, excluding GO involved in transcription or cell cycle regulation all the remaining items were directly associated with the development (Fig. 4E) confirming the strong downregulation bias in the expression of developmental genes initially observed with the analysis of the SR TFs.

**Fig 4.**
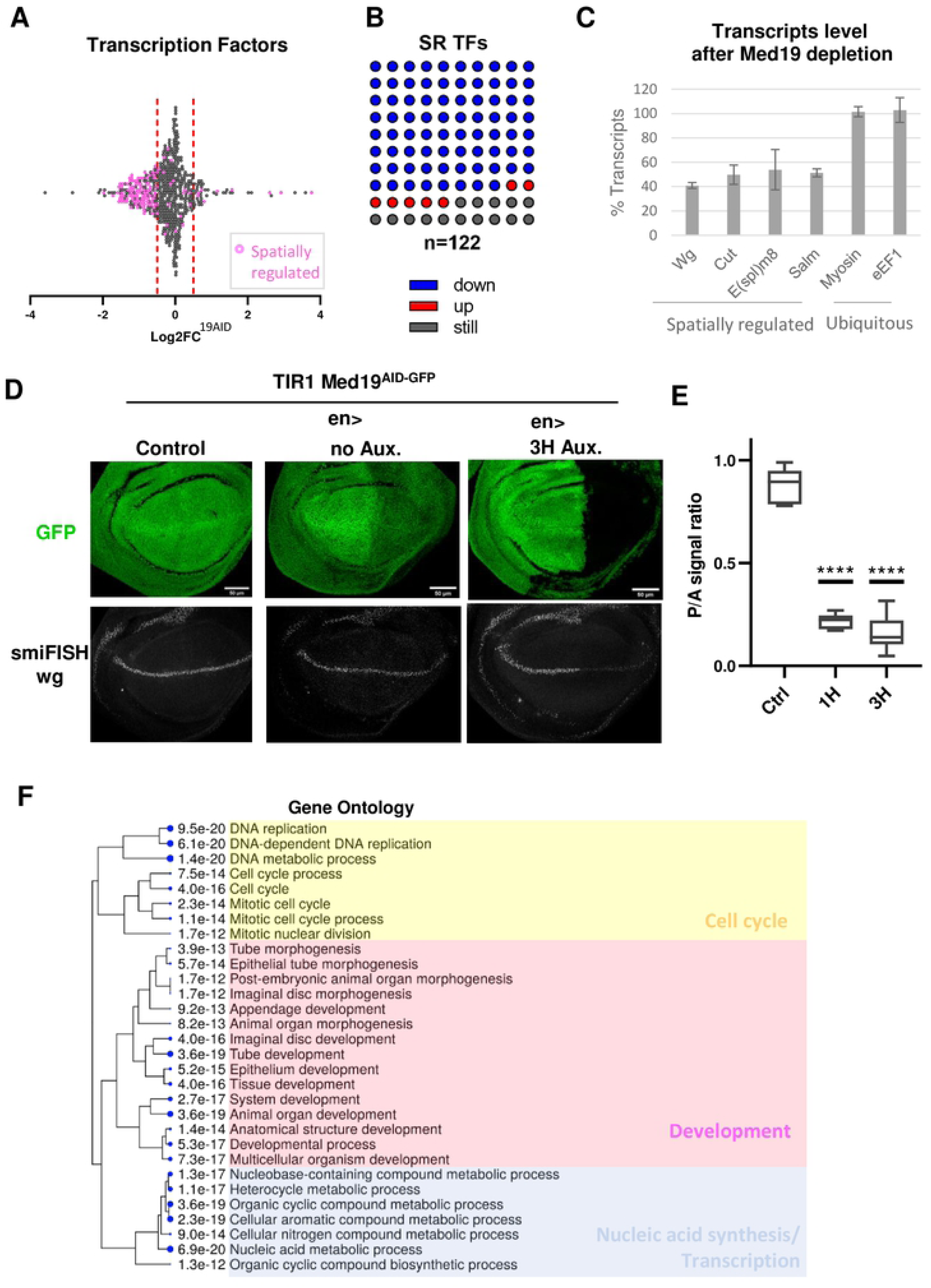
Med19 promotes the expression of the spatially regulated developmental genes. (A) Scatter plot depicting the distribution of the Log2FC values of the 573 transcription factors found significantly expressed in the wing imaginal disc from the RNAseq data. The transcription factors with a demonstrated spatial transcriptional regulation (SR TFs) in the wing imaginal disc are shown in purple (n=122). (B) Dot plot 10x10 showing the proportion of the non-regulated (dark grey), upregulated (red), or downregulated (blue) genes among the 122 SR transcription factors. (C) Bar graph made from RT-qPCR data showing the relative level of transcripts of 6 genes known to be spatially regulated (*wg*, *cut*, *E(Spl)m8*, *sal* or ubiquitous (Myosin, E2F1) in the wing disc, after Med19^AID^ depletion was carried out for 4 hours in all the disc cells (percentage of the control in which Med19^AID^ degradation was not triggered). The graph shows the mean and standard deviation of 3 independent experiments. (D) Confocal images of the wing pouch region of dissected *en>; Med19^AID^* wing discs that were subjected to Med19AID degradation (en> 3H Aux), or not (control: no Gal4 driver, and *en*> no Aux), and in which *wingless* (wg) pre-mRNA were visualized by smiFISH using intronic probes. top images: Med19^AID^ GFP signal. bottom images: *wg* smiFISH signal. (E) Wing disc D/V boundary posterior to anterior wg smiFISH signal ratio established in *en>; Med19^AID^* wing discs: no auxin control (no Aux. n=5), 1H auxin (n=6), and 3H auxin (n=6). ****: P value <0.0001 (Unpaired t test). (F) Hierarchical clustering tree of the 30 best FDR of the GO terms associated with the downregulated genes. The size of the solid circles is proportional to the enrichment False Discovery Rate (FDR). GO clusters appear related to development (purple), nucleic acid synthesis (blue), and cell cycle regulation (yellow).

### Regulated expression of Notch responsive *E(spl)-C* genes requires Med19 activity

The spatio-temporal control of the developmental gene expression is achieved through the action of multiple signaling pathways, among which Notch. *Wg* expression at the wing D/V boundary is a well-established marker of Notch (N) activity [30], raising the idea that Med19 could function as a transcriptional effector of the Notch signaling pathway, an hypothesis we further addressed experimentally. Notch, a key player in the control of wing development, regulates the expression of a large number of genes in the wing imaginal disc, among which the eleven genes of the *Enhancer of Split* complex (*E(spl)-C*) that are directly activated by the Notch intracellular-Domain (NICD) / Suppressor of Hairless (Su(H) CSL in mammals) complex [31]. Remarkably, the RNA-seq DGE analysis indicated that the majority of the *E(spl)-C* genes were downregulated upon Med19^AID^ degradation (Figure 5A). RT-qPCR analysis confirmed that the expression of *E(spl)m8* is compromised after Med19^AID^ removal in the wing imaginal disc (Fig. 4B). In addition, wing disc *in situ* hybridization was performed to follow the expression of two other genes of the *E(spl)-C*, m3 and mβ, which display well-characterized expression patterns in the wing pouch [32]. RNA FISH revealed that the transcription of both *E(spl)* genes was severely reduced specifically in the region of the disc where the depletion of Med19^AID^ was specified (Fig. 5 B and C) therefore confirming the RNA-seq data. The fact that the expression of the known transcriptional regulators of the E(spl)-C (including Su(H)) are not affected after Med19^AID^ depletion (Fig. 5A), and that the Mediator Complex and Su(H) have been previously reported to physically interact [33] raised the possibility that Med19 could function as a cofactor of Su(H) in the expression of *E(spl)-C* genes. We experimentally addressed this idea by examining whether Med19 and Su(H) could directly interact *in vitro*. We found that bacterially produced GST-Med19 specifically pulled down Su(H) molecules synthetized *in vitro* (Figure. 5D), providing evidence that both proteins associate with each other.

**Fig 5.**
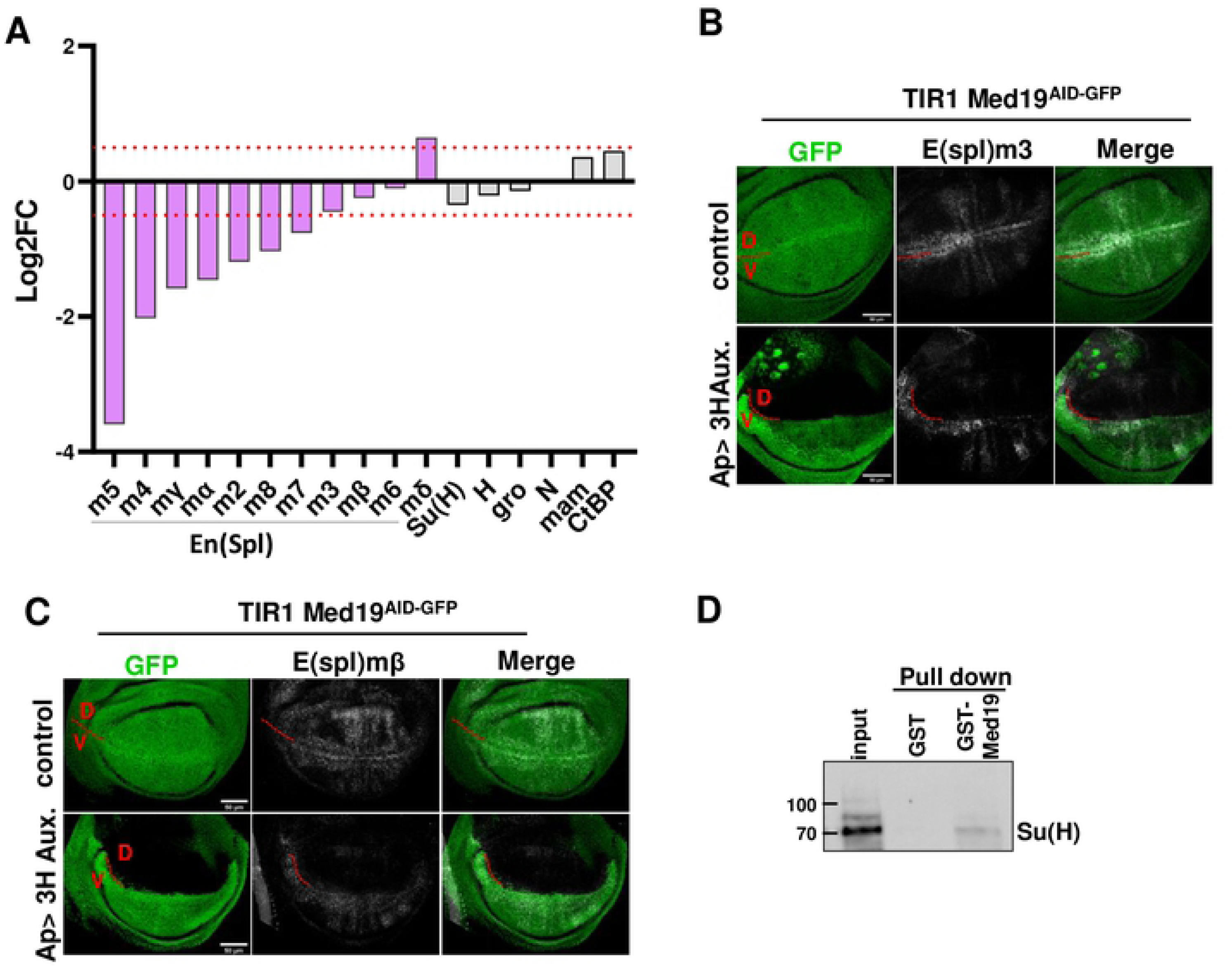
Med19 is required for the expression of Notch responsive genes in the wing imaginal disc. (A) Scatter plot and median of the expression fold change (Log2FC), obtained from the Med19^AID^ RNAseq DGE analysis, of the 11 Enhancer of Split complex (E(spl)-C) genes, and genes encoding TFs involved in the transcriptional regulation of the E(spl)-C genes, namely Suppressor of Hairless Su(H), Hairless (H), Groucho (Gro), Mastermind (Mam), Notch (N), and CtBP. The red dotted lines delineate the -0.5<Log2FC<0.5 expression fold change zone. (B, C) Maximum projection confocal microscopy images of Med19^AID^ GFP signal and transcripts detection using smiFISH probes in the pouch of wing imaginal discs in which Med19^AID^ has been regionally degraded, or not. Driving the degradation of Med19^AID^ for 3 hours in the dorsal domain of the wing disc using the *ap*>; Med19^AID^ genetic set-up severely affects the level of *E(spl)m3* (B), and *E(spl)mbeta* (C) transcripts specifically in the area where Med19 is depleted. (D) Western blot analysis of a GST pull-down experiment showing that *in vitro* translated HA-tagged Su(H) – detected using an anti-HA antibody – physically interacts with GST-Med19, but not with GST. Input: 5% of the starting material.

**Fig 6.**
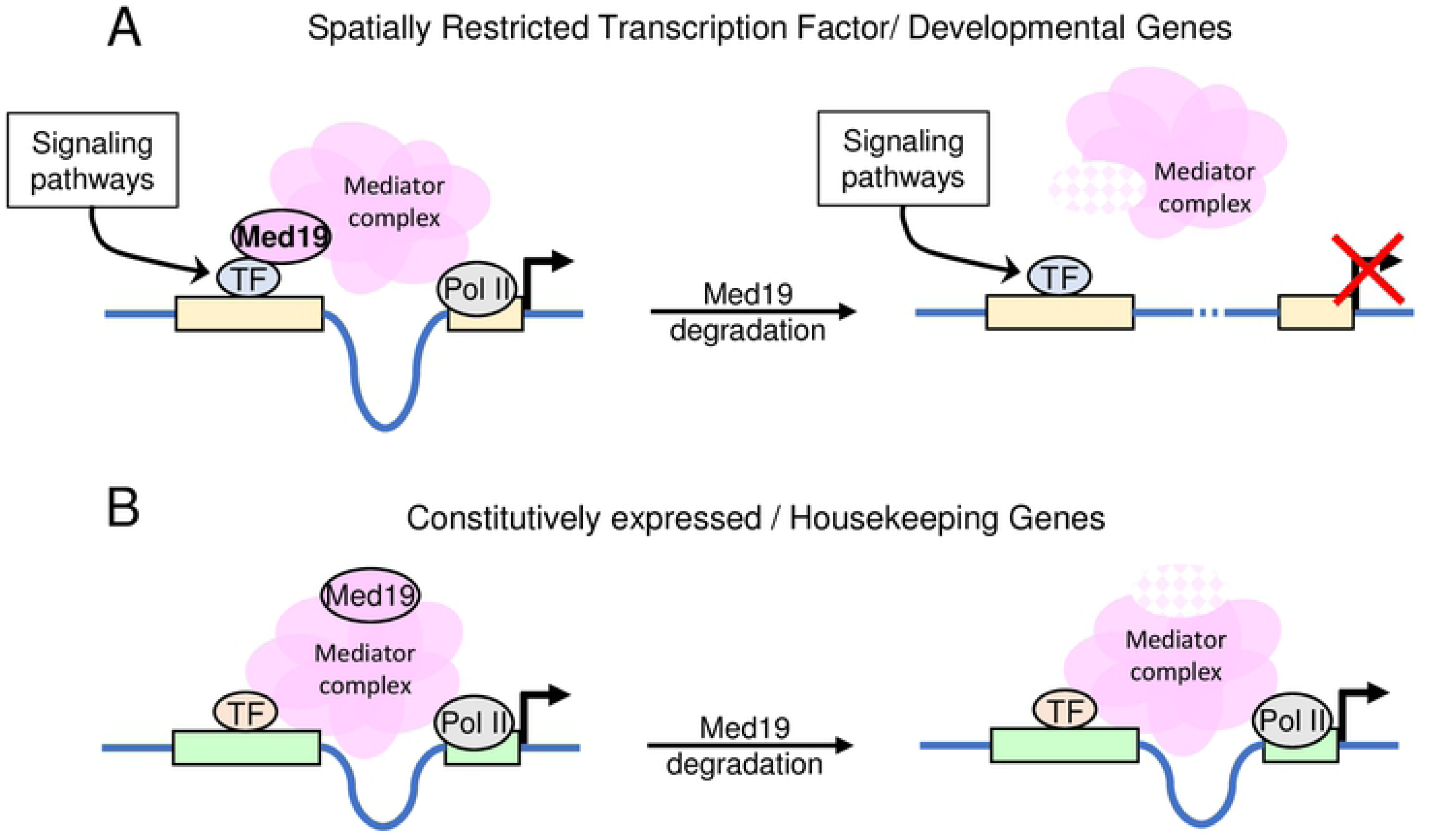
Model, derived from the analysis of the nature of the genes deregulated or not after Med19^AID^ degradation, showing how Med19 may function. (A) Med19 acts as a critical cofactor of transcription factors whose activity is regulated by signaling pathways, in negative or positive control of transcription. (B) Med19 is not a Mediator subunit required for the expression of the constitutively expressed genes.

Taken together, these data support a mechanistic model where Med19 promotes the expression of the Notch responsive *E(spl)-C* genes through a direct interaction with the transcription factor Su(H).

## Discussion

The role of the Mediator subunit Med19 on gene expression remains poorly documented due to the limited number of studies that analyzed the consequences of its loss of function on the transcriptome [6,34,35]. The present study addresses this question in a comprehensive way, in the context of the development using the *Drosophila* wing imaginal disc as an *in vivo* developmental model. Our project to identify the genes directly controlled by Med19 in this developing tissue faced the issue of the deleterious effect of losing *Med19* function over long period of time. The problem was that conventional loss-of-function strategies such as the use of a null allele or RNA interference, provide poor temporal control and would have severely compromised the obtention of wing imaginal discs in which complete depletion of Med19 in all the cells of the tissue occurred. On top of that, prolonged depletion causes distortion of the gene expression profiling data due to transcriptional cascade effects and stress responses, jeopardizing the recognition of the genes directly regulated by Med19 activity. In this context, the strength of our study is the use of the AID targeted protein degradation technology to generate genetically modified flies in which rapid proteolysis of the endogenous Med19 can be triggered in the tissues of the living animal upon request. This unique tool enabled to establish the gene expression profiling in wing imaginal discs in which Med19 ablation occurred for a defined short period of time, and therefore we have been able to get access to the early consequences of losing *Med19* function on gene expression.

Overall, the analysis of the expression change data reveals that Med19 up and down-regulates less than a quarter of the genes expressed in the wing imaginal disc, therefore supporting the notion that this Mediator subunit carries out specific functions. Although the Med19 KO studies done in mouse reflect late consequences of Med19 loss, the restricted and bidirectional nature of the gene deregulation has also been described, in line with our results [6, 35], and beyond *Med19*, this trait appears to be a common phenotype associated with the loss of function of several Mediator subunits .

It is currently established that the transcription of the vast majority of the RNA PolII-dependent genes requires the Mediator complex. Therefore, the fact that the expression of a large number of genes is not affected, or even upregulated after Med19 depletion implies that the Mediator complex devoid of Med19 keeps most of its functionality. This conclusion is supported by the fact that in yeast, a Mediator complex missing Med19 can be biochemically isolated [36] and that such incomplete, but still structured, complex has been visualized by cryo-electron microscopy [37].

By crossing our expression change results with expression profile data, we discovered the existence of a link between the dependence of a gene on Med19 for its expression and its mode of expression: constitutive versus highly regulated. More precisely, we found that Med19 is primarily involved in the control of the expression of dynamically regulated genes (referred here to as SR genes). However, the most unexpected result of this study is the fact that the vast majority of the constitutively expressed genes (referred here to as HK genes), among which the housekeeping genes, do not rely on Med19 for their expression. At least, this finding is in agreement with the fact that Med19 is not strictly required for cell viability as shown by studies made in *Drosophila* and Mammalian cells [6, 14], and supports the notion that this Mediator component does not exert essential functions in the non-regulated – constitutive – mode of expression, which represents about half of the expressed genes.

It is likely that our observations are the reflect of the fact that SR and HK use separate mechanisms of expression. Indeed, it is established that SR and HK genes differ in their genomic organization [38], their enhancer and core promoter structure and composition, and each class of genes exhibit mutually exclusive enhancer-core promoter functional relationships [39, 40]. In line with our findings pointing to a specific role of Med19 in the regulated, but not in the constitutive mode of gene expression, it has been recently shown that some Mediator subunits display a marked functional preference for the core promoter of genes expressed highly variably and having developmental functions, but are unable to function with core promoters of HK genes [41]. Furthermore, genome-wide binding studies revealed that the Mediator complex appears preferentially concentrated to a special type of chromatin functioning as transcriptional regulatory hubs, referred to as super enhancers in mammals, controlling the expression of dynamically regulated genes [42, 43].

In a context where the expression change after Med19 loss is globally bidirectional, we have been able to identify a functional class of genes, the spatially regulated transcription factors, and more generally the developmental genes as revealed by the gene ontology analysis, whose expression requires Med19. Although the existence of super enhancers has not been formally proven in *Drosophila*, this observation could be related to the finding that in mammalian cells the Mediator plays a critical role at this special class of enhancers to promote the expression of lineage-specifying genes, and especially those encoding the transcription factors responsible for the cell identity [7].

More investigations will be required to understand the reasons of the involvement of Med19 in promoting expression of the developmental genes. Specifically, approaches allowing to quantitatively map the interaction between Mediator and RNA PolII subunits and the genome, such as ChIP-seq, may provide clues to key questions like whether the Mediator complex is still associated to the enhancers of developmental genes after Med19 ablation, the expression collapse is linked to a defect in the pre-initiation complex (PIC) formation, or the problem arises from a defect in the transcriptional pausing release.

We experimentally confirmed the finding, made from the RNA-seq analysis, that Med19 is important for the transcriptional activation of highly regulated developmental genes by showing *in situ* that some canonic target genes of the Notch pathway in the wing imaginal disc need Med19 activity for their expression. In particular, the visualization of the drop in the level of *wg* pre-mRNA shortly after Med19 degradation demonstrates the existence of consequences at the level of the transcription, therefore excluding the possibility that our observations are exclusively linked to a post-transcriptional effect such as transcripts degradation. We and others have shown that Med19 acts as a cofactor of transcription factors with which it interacts, these including in *Drosophila* the HOX [14], GATAs [16], and the corepressor Brakeless [44]. Although further work is needed to characterize the interaction between Med19 and the transcriptional regulator Su(H), we propose that Med19 works in combination with Su(H) in the transcriptional activation of the Notch target genes.

In conclusion, this work points the idea that finely regulated modes of transcription that use complex molecular mechanisms may require the functionality of all the subunits of the Mediator complex whereas mechanistically simpler constitutive expression may only rely on a subset of the Mediator components. To get insight into this notion, the experimental approach we developed could be extended to each of the 30 Mediator subunits. Such a comprehensive study will enable to know to which extent the majority of the Mediator subunits, like Med19, specifically operate regulated genes. Critically, the knowledge of the transcriptional landscape regulated by each of the Mediator components in a given biological context will be essential to our understanding of how the Mediator operates.

## Acknowledgements

We are grateful to the genotoul bioinformatics platform Toulouse Midi-Pyrenees and Sigenae group for providing help and storage resources on the local Galaxy instance http://sigenae-workbench.toulouse.inra.fr. We thank M. Aguirrebengoa from the big-A facility of CBI and Alexandra Manchego for their help in bioinformatics and statistical analyses, J. Favier for the CBI *Drosophila* genome edition facility, F. Payre, and A. Vincent for critical reading of the manuscript. We acknowledge the Bloomington *Drosophila* Stock Center for providing transgenic flies. Lastly, we thank the Toulouse RIO Imaging platform.

## Supporting information

**S1 Fig.** Scheme depicting the CRISPR/Cas9-based strategy used to generate *Med19^AID^*, a *Med19* degradable and fluorescent allele. The donner construct (middle) highlights the nature of the homology arms used to introduce the AID-GFP cassette by homologous recombination. Position of the PCR primers used to characterize the wt vs 19AID allele is shown.

**S2 Fig. Degradation of Med19^AID^ in the posterior compartment of the wing imaginal disc using the *engrailed* driver (*en*>)**. (A) Confocal images of wing discs dissected from *en>TIR1 Med19^AID^* larvae either fed for one hour with NAA (1H), or mock treated (no auxin). (B) Box plot showing the Med19AID depletion level calculated as the posterior to anterior GFP intensity ratio in the wing pouch of *en>TIR1; Med19^AID^* wing imaginal discs (n=5) obtained and imaged as in (A).

**S3 Fig. The expression of most of canonical housekeeping (HK) genes is not affected by Med19^AID^ depletion.** Scatter plot analysis of the expression fold change (Log2FC) of genes belonging to several families of housekeeping functions (x axis, list detailed in table S3). The red dashed lines mark Log2FC= +/ - 0.5.

**S4 Fig. Venn diagram showing the overlap between HK - HK* (top), and SR - SR*(bottom) lists of genes.**

**S5 Fig. The expression of *achaete* (*ac*) and *wingless* (*wg*) is impaired specifically in the domain of the wing imaginal disc where Med19^AID^ is degraded.** (A) Confocal images of *Ap>TIR1; Med19AID* wing imaginal disc immunostained for Ac and GFP (Med19^AID^ detection). Upper row: control wing disc in which Med19^AID^ degradation did not occur (no driver). Bottom row: wing disc in which Med19 degradation was maintained for 3 hours in the dorsal compartment, using an *apterous* driver. Right column: magnification in the anterior wing pouch Ac staining pattern emphasizing the loss of Ac expression in the dorsal row of cells of the anterior wing margin when Med19^AID^ degradation is triggered in the dorsal compartment of the wing disc. (B) *en>TIR1; Med19AID* wing imaginal discs immunostained for Wg and GFP (Med19^AID^), in which the degradation of Med19^AID^ was operated with the *en*> driver for 3 hours (bottom row), or not triggered (control, upper row).

**S1 Table**: Spreadsheet of the sequence of all the DNA oligonucleotides used in this study.

**S2 Table**: List of the 58 *Drosophila* tissues and developmental stages (modENCODE project) used to establish the RNA expression profile of the genes expressed in the wing imaginal disc.

**S3 Table**: List of canonical housekeeping genes.

**S4 Table**: Spreadsheet containing the lists of HK and SR genes (published), and the HK* and SR* genes.

**S5 Table**: 30 best GO terms of the down-regulated genes sorted by FDR enrichment obtained on ShinyGO web site.

